# Colonization of naïve roots from *Populus tremula x alba* involves successive waves of fungi and bacteria with different trophic abilities

**DOI:** 10.1101/2020.11.04.368126

**Authors:** F. Fracchia, L. Mangeot-Peter, L. Jacquot, F. Martin, C. Veneault-Fourrey, A. Deveau

**Author notes:** F.F and L.M-P contributed equally to this work. C.V-F and A.D contributed equally to this work. Corresponding author Mailing address: UMR1136 INRAE Université de Lorraine, Interactions Arbres Micro-organismes, 54280 Champenoux, France.

## Abstract

Through their roots, trees interact with a highly complex community of microorganisms belonging to various trophic guilds and contributing to tree nutrition, development and protection against stresses. Tree roots select for specific microbial species from the bulk soil communities. The root microbiome formation is a dynamic process but little is known on how the different microorganisms colonize the roots and how the selection occurs. To decipher if the final composition of the root microbiome is the product of several waves of colonization by different guilds of microorganisms, we planted sterile rooted cuttings of Gray Poplar obtained from plantlets propagated in axenic conditions in natural soil taken from a poplar stand. We analyzed the root microbiome at different time points between 2 and 50 days of culture by combining high-throughput Illumina MiSeq sequencing of fungal rDNA ITS and bacterial 16S rRNA amplicons with Confocal Laser Scanning Microscope observations. The microbial colonisation of poplar roots took place in three stages but the dynamic was different between bacteria and fungi. Root bacterial communities were clearly different from the soil after two days of culture. By contrast, if fungi were also already colonizing roots after two days, the initial communities were very close to the one of the soil and were dominated by saprotrophs. Those were slowly replaced by endophytes and ectomycorhizal fungi. The replacement of the most abundant fungal and bacterial community members observed in poplar roots along time suggest potential competition effect between microorganisms and/or a selection by the host.

**Importance:** The tree root microbiome is composed of a very diverse set of bacterial and fungal communities. These microorganisms have a profound impact on tree growth, development and protection against different types of stress. They mainly originate from the bulk soil and colonize the root system which provides a unique nutrient rich-environment for a diverse assemblage of microbial communities. In order to better understand how the tree root microbiome is shaped along time, we observed the composition of root-associated microbial communities of naïve plantlets of poplar transferred in natural soil. The composition of the final root microbiome rely on a series of colonization stages characterized by the dominance of different fungal guilds and bacterial community members along time. Our observations suggest an early stabilization of bacterial communities, whereas fungal communities are established following a more gradual pattern.

## Introduction

Trees have been recognized as metaorganisms possessing specific microbiomes that are key determinants of tree health and productivity (1). The tree root microbiome, notably its fungi and bacteria, is particularly important as it participates to nutrients and water acquisition and in the protection of trees against pathogens (1; 2). Despite the overall positive effects of microbiome on their hosts, the different members of the microbiome can have contrasted effects: some such as mycorrhizal symbionts can be beneficial by promoting plant nutrition and resistance against stresses, while others such as pathogens are detrimental (3; 4; 5). In addition, species of beneficial microorganisms may differ in their activities although redundancy between species exist (6). For instance, some bacterial species promote the growth of their hosts by producing phytohormones that stimulate the growth of the root systems while other facilitate the access to key nutrients (7). Similarly, ectomycorrhizal fungi (EcM) that provide nitrogen, phosphorus and oligonutrients in exchange of carbon, can strongly differ among species and strains in their abilities to access key nutrients in soil (8). If the main abiotic (e.g. edaphic properties, climate) and biotic (e.g. genotype, root exudates) factors that influence the composition of the root microbiome are now well documented (9; 10), little is known on how occur the assembly of the tree root microbiome.

Roots are mainly colonized by microorganisms found in the surrounding soil that serve as a seed bank. Colonization occur in a two-step process in which root exudates initiate the recruitment in the rhizosphere - the soil area directly under the influence of plant roots -, followed by the entry inside the root tissues and a fine tuning of the communities of the rhizoplane – the roots surface – by the plant-microbial interactions (9; 11). Root exudate chemistry and dynamic, together with microbial preferences for substrates, determine the assembly of bacterial community of the rhizosphere in some annual plants (12; 13). However, by comparison, the tree root microbiome is much more complex as it harbours on top of highly diverse bacterial communities, a plethora of microorganisms with potential different functional capacities compared to herbaceous plants. Indeed, tree roots are colonized by EcM in temperate and boreal forest ecosystems (1) but also by endophytes and saprotrophic fungi even though their role still remains elusive (14; 15). In addition, some trees of temperate climates such as maple and alder trees associate with arbuscular mycorrhizal fungi (AM). Finally, a few trees like poplars and eucalypts are colonized by both EcM and AM. While some of these microorganisms can react to root exudates, others rely on specific molecular dialogues with their hosts to establish themselves in the root system (16). In addition, bacteria and fungi colonizing tree roots are susceptible to interact together through faciliation (17; 18) and/or competition events (19). Finally, trees are long-lived woody perennial plants with a different management of the nutrient allocation compared to herbaceous and annual plant species such *Arabidopsis thaliana* or crops (20; 21).

The establishment of the root microbiome is a dynamic process where specific microbial communities progressively colonise root systems under both the plant selection and the interactions among microorganisms. Previous works on root colonization have been carried out on trees to understand the mechanisms of the establishment of the tree root microbiome and of tree root selection. For instance, aspen root colonization by the plant growth promoting bacteria *Pseudomonas* indicated that the spatial and temporal patterns of colonization of roots was different between the four strains of bacteria and was correlated with the ability of bacteria to form biofilm (22). In pine roots, comparison of the dynamic of root colonization of two EcM fungi revealed different strategies. The ability of *Rhizopogon* to colonize roots rapidly from spores and its important early abundance constrasted with later root colonization and the slow increase in abundance of *Tomentella* (23). The work of (24) and (25) on EcM and AM colonisation dynamic in eucalyptus roots showed a successionnal replacement of AM by EcM fungi. Similarly, (26) showed negative associations among EcM and AM fungi leading to a depletion of AM and an increase in EcM in lateral roots in poplar. Nevertheless, these studies focused on one or few bacterial or fungal species using microbial inoculation and did not look at the overall dynamic of the microbiome, including endophytes and saprophytes. Yet, pioneering studies on ectomycorrhizal and bacterial communities of the roots of Pines indicate that the full microbiome is likely subjected to a complex dynamic during the colonization process (27). Investigating the temporal succession of microbial communities colonizing root system of young naïve trees is needed to help the understanding of the complex interactions occurring between microbiota and their host trees. *Populus* is a good model to address such question because it is now a well-established model to study tree microbiome (15; 28; 29; 30; 31) and it hosts both EcM and AM, fungal endophytes and bacterial communities (32; 33). In addition, poplar clones can be cultivated *in vitro* in sterile conditions, thus limiting genetic variability and allowing to focus solely on the colonization by communities coming from the soil and not vertically transmitted. Last but not least, poplar is an important species in the Northern hemisphere forestry with 80 million hectares of trees in the world (FAO, 2004). In France, poplar culture represent 23% of the annual broad leaves trees yields and french industries should have difficulties in supply in 2023 (source CODIFAB).

Based on these previous studies, we hypothesized that fungal and bacterial communities originating from the natural soil successively colonized host roots with a progressive replacement of the root microbiota members. To test this hypothesis, we used the Gray poplar, *Populus tremula x alba* as a woody and perennial model organism and we assessed the dynamic of tree roots colonization by fungal and bacterial communities during the first 50 days of contact between naive tree roots and soil microbial communities by 16S and ITS rRNA gene-targeted Illumina MiSeq sequencing and Confocal Laser Scanning Microscopy (CLSM).

## Results

### Plantlet development and ectomycorrhiza formation in natural soil

In order to investigate the temporal colonisation dynamic of *Populus* roots by fungal and bacterial communities, three-week-old axenic cuttings of poplar were planted in pots containing natural soil taken from a poplar plantation (**Figure S1**). Monitoring of the growth of the root systems indicated a slow development of the roots during the first 15 days followed by an increased growth in the next weeks (**Figure 1 A**). First short roots and ectomycorrhizae were observed at 10 and 15 days (T15), respectively. The rate of ectomycorrhiza formation regularly increased to reach 37% at 50 days (T50) post plantation and nearly doubled between T15 and T50 (**Figure 1 B**).

**Figure 1.**
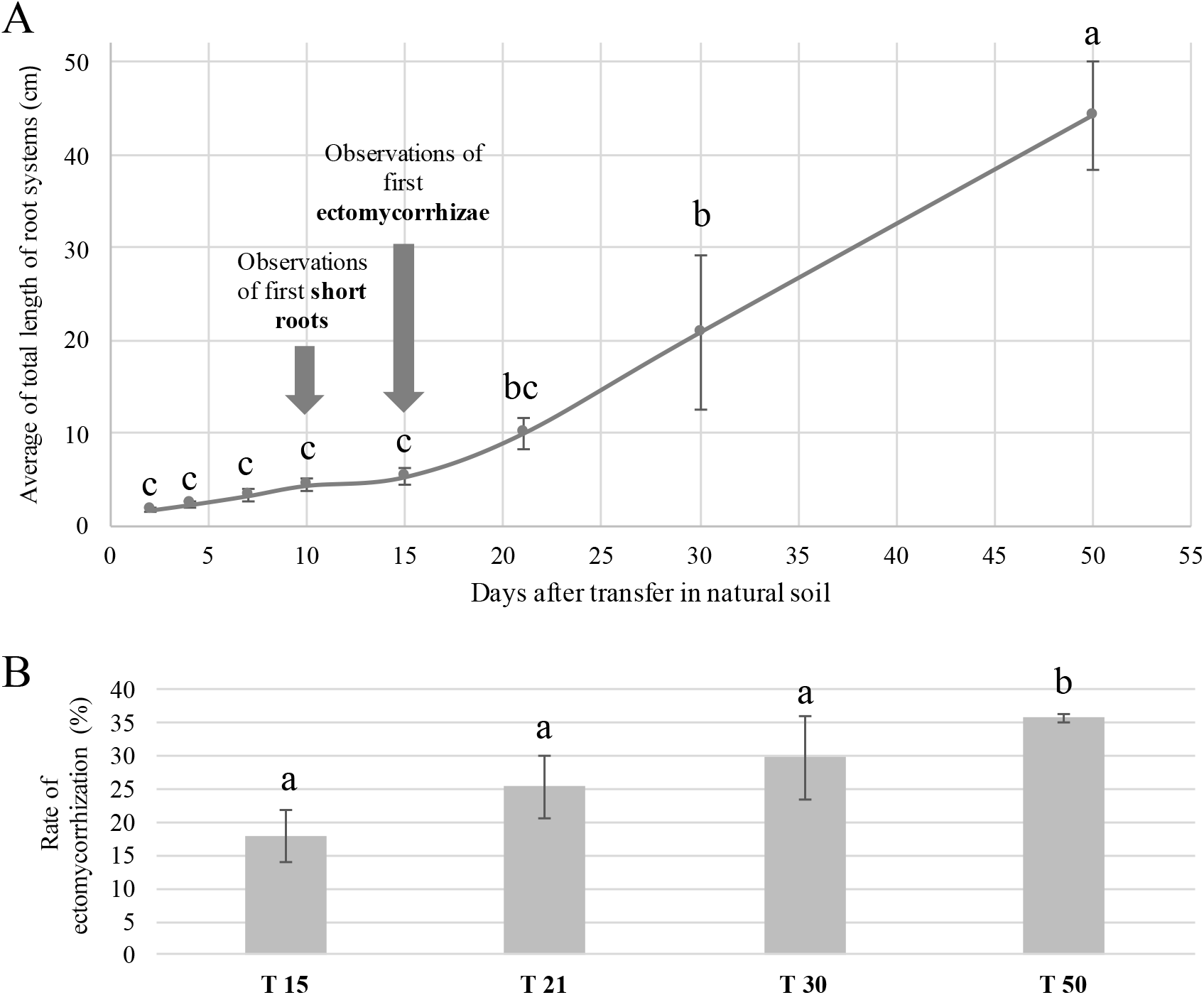
Root development and ectomycorrhizae formation along time. **A**. Total length of the root system measured at each sampling time from T2 to T50. **B**. Ectomycorrhization rate of Populus roots from T15 to T50 calculated as the number of fungal colonized lateral roots/ total number of lateral roots. Each given value is the average value of 7 replicates +/− SE. Different letters denote significant difference between each sampling time (One-way ANOVA, factor=sampling time, P<0.05).

Fungal and bacterial colonization of the roots were tracked using two complementary methods: 16S and ITS rRNA gene-targeted Illumina MiSeq sequencing and confocal microscopy. No amplification of ITS and 16S rDNA genes were obtained from samples of roots collected before vitroplants were transfered in natural soil. These results are in accordance with CLSM observations concerning fungal colonisation as no fungal structure could be visualised at T0 (**Figure S2**). These results validate the axenic status of the in vitro root systems.

### Microbial sequencing

Sequencing of ITS2 and 16S rDNA amplicons were performed on bulk soil and roots DNA samples between T2 and T50. After quality filtering and chimera removal, a total of 909,764 fungal reads and 1,678,387 bacterial reads were kept for further analyses. After taxonomic assignment, elimination of contaminants and completion of normalization by rarefaction, 373 fungal OTUs (70 ± 1 per sample) and 887 bacterial OTUs (518 ± 6 per sample) were detected. One of the five biological replicates of T4, T10 and T15 and one of the five biological replicates of T2 and T50 were eliminated from the data after completion of rarefactions of ITS and 16S data, respectively.

### Soil microbiome composition

Since bulk soil is the only reservoir of microorganisms for the colonisation of the root system in our conditions, we first analysed its composition at T0. It was heavily colonized by complex fungal and bacterial communities, as expected from a previous study on soil taken from the same poplar plantation (48). Twenty bacterial phyla, 111 families and 633 OTUs were detected. The community was dominated by three phyla: Acidobacteriota (27.8 ± 0.2% of the relative abundance), Verrucomicrobiota (26.8 ± 1.5%) and Proteobacteria (23.1 ± 0.4%, **Table S1**). OTUs from genus *Candidatus Udeobacter* accounted for 22% (± 1.4) of the relative abundance by themselves (**Table S1**).

Regarding fungi, Basidiomycota represented the most abundant phylum (45.3 ± 3.1%), followed by Ascomycota (31.0 ± 3.2%) and Mucoromycota (11.7 ± 0.9%, **Table S2**). At a lower taxonomic scale, 102 fungal species and 156 OTUs were detected in the bulk soil. *Sebacina* was the most dominant and represented up to 17% (± 0.5) of the relative abundance, followed by *Umbelopsis dimorpha* (10.7 ± 0.8%) and *Mortierella* (6.1 ± 0.5%, **Table S2**). Twenty EcM (34.2 ± 3.1% of the relative abundance), 31 saprotrophs (20.3 ± 2.5%) and 6 potential endophytes species (9.0 ± 1.1%, **Table S3**) were found.

### Overall dynamic of microbial colonization of the root system

First bacterial and fungal colonisations of *Populus* roots were observed after two days of growth (T2). The number of bacterial and fungal OTUs detected in roots was 1.3 and 2.1-fold lower at T2 than in the bulk soil (**Table 1**, Kruskal-Wallis, P.adj <0.05). Then, diversity of bacterial and fungal communities colonizing the roots evolved differently. Bacterial richness and diversity measured by Shannon index significantly increased over time to stabilize around 21 days. By contrast, fungal richness reached a maximum at 15 days then decreased, while fungal diversity slowly decreased from T15 to T50 (**Table 1**).

**Table 1.** Diversity of the bacterial and fungal communities detected in soil and in roots across time. Richness and Shannon indexes calculated for bulk soil and root samples collected at the different sampling time from T2 to T50. Each given value is the average value of 4 or 5 replicates +/− SE. The asterisks denote significant difference between bulk soil and roots collected at T2 and different letters denote significant differences between each sampling time from T2 to T50 (Kruskal-Wallis, correction Bonferroni, Fisher’s LSD post-hoc test, P.adj<0.05).

| Sampling time | Bacterial richness | Bacterial shannon index | Fungal richness | Fungal shannon index |
| --- | --- | --- | --- | --- |
| Bulk soil | 620.3 ± 6.3 * | 5.30 ± 0.05 * | 155.0 ± 1.0 * | 3.7 ± 0.1 * |
| T2 | 463.0 ± 31.3 c | 3.2 ± 0.3 a | 66.0 ± 7.8 abcd | 2.5 ± 0.1 |
| T4 | 414.8 ± 54.0 c | 3.5 ± 0.2 ab | 63.3 ± 7.5 bcd | 2.4 ± 0.2 |
| T7 | 494.0 ± 35.1 bc | 3.7 ± 0.2 abc | 74.4 ± 8.3 abc | 2.7 ± 0.3 |
| T10 | 437.4 ± 35.8 c | 3.3 ± 0.3 abc | 68.8 ± 11.7 bcd | 2.3 ± 0.2 |
| T15 | 464.0 ± 48.0 c | 4.0 ± 0.2 bc | 89.8 ± 4.6 a | 2.4 ± 0.3 |
| T21 | 616.8 ± 18.6 a | 3.9 ± 0.4 bc | 87.6 ± 5.3 ab | 2.3 ± 0.1 |
| T30 | 598.0 ± 46.7 ab | 4.4 ± 0.2 c | 58.8 ± 3.5 cd | 1.7 ± 0.1 |
| T50 | 648.6 ± 11.0 a | 4.8 ± 0.1 c | 50.8 ± 3.8 d | 1.5 ± 0.2 |

These dynamic changes of richness and diversity were associated with modifications of the structure of the microbial communities between bulk soil, suggesting an early selection of root microbial communities (**Figure 2**, PERMANOVA, P.adj <0.05, Time explained 45% of the variance for bacteria and 38% for fungi). The structures of root bacterial and fungal communities then progressively evolved from T2 to T50 although close time points were not statistically different (e.g T2-T4, T15-T21…, pairwise PERMANOVA, P.adj <0.05; **Figure 2**). Three stages of bacterial and fungal colonization were defined based on NMDS graphic representations and pairwise PERMANOVA: an “early” stage from T2 to T4, one “Intermediate” from T7 to T15 and one “Late” from T21 to T50.

**Figure 2.**
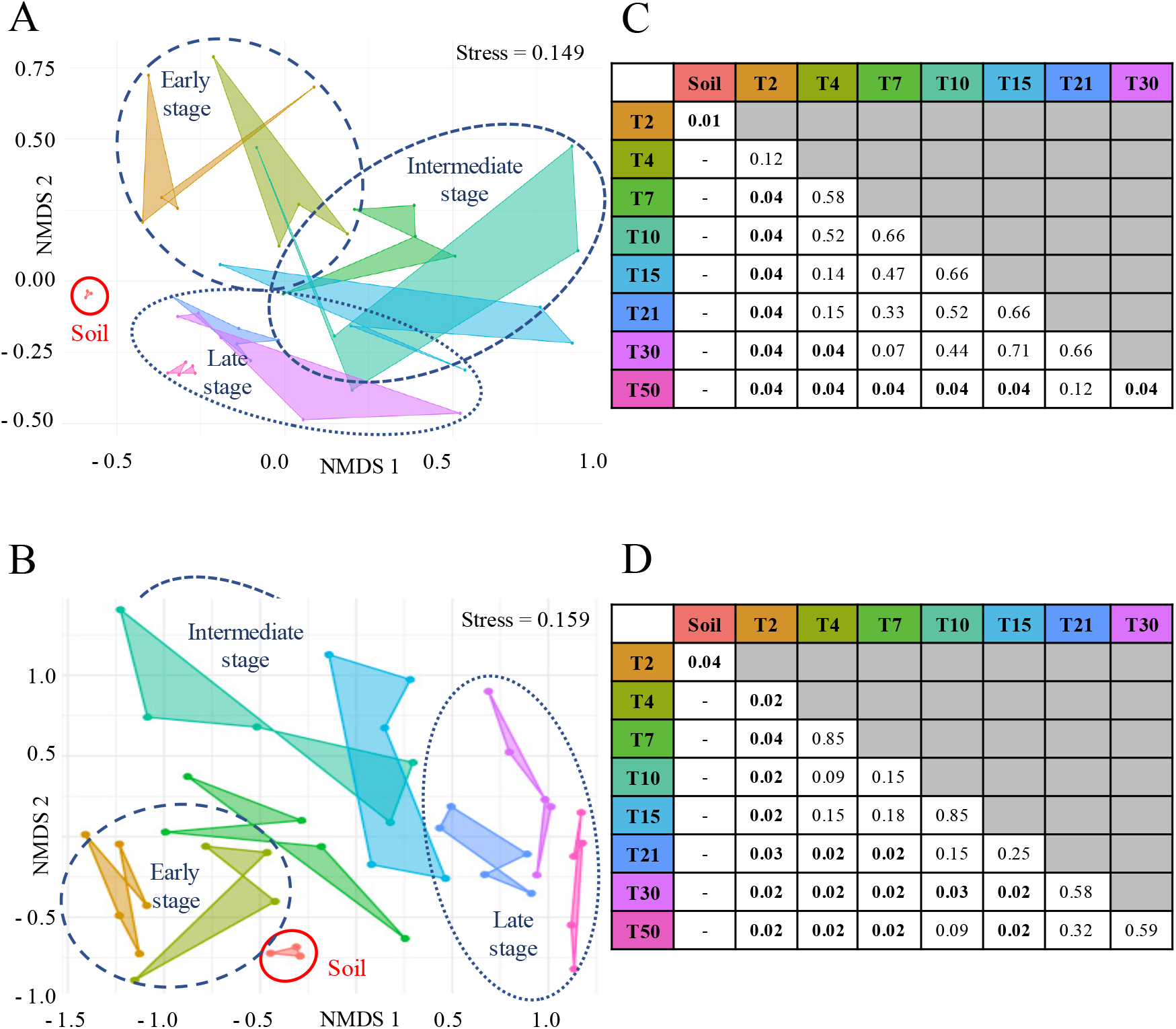
Structure of bacterial and fungal communities colonizing Populus roots across time. Non-metric multidimensional scaling (NMDS) ordinations of bacterial OTU (**A**) and fungal species (**B**) across compartments (bulk soil and roots) and sampling times (from T2 to T50) based on Jaccard distance. Adjusted P-values of variances explanation based on pairwise comparisons using perMANOVAs on the Binary distance for bacterial OTU (**C**) and fungal species (**D**).

### Assembly of fungal and bacterial communities in the roots at the early stage

The composition of both bacterial and fungal communities already strongly differed from the one of the bulk soil at T2 although this effect was much more pronounced for bacterial communities (**Figure 3**). Fifteen out of the 20 bacterial phyla detected in the bulk soil showed significant differences of their relative abundance in roots at T2 (**Table S1**, Kruskal-Wallis, P.adj < 0.05). Most striking was the massive colonization of the roots by Proteobacteria which accounted for 79% of all phyla, while they were 3.5-fold less present in the bulk soil (**Table S1**, P.adj = 0.034). By contrast, Acidobacteria and Verrucomicrobia were poorly represented in roots at T2 while they dominated in the bulk soil (**Table S1**). Out of the 157 genera detected in the roots at T2, the bacterial communities were dominated by 7 genera that all together accounted for 56% of the relative abundance. Among those, four were members of the *Burkholderiaceae* family and the genus *Burkholderia* accounted by itself for 35% (± 5) of the relative abundance. Members of *Mucilagenobacter* (Sphingobacteria), *Pseudomonas* (γ-Proteobacteria) and *Streptomyces* (Actinobacteria) were also quite abundant as their relative abundance exceeded 3% at this early time point.

**Figure 3.**
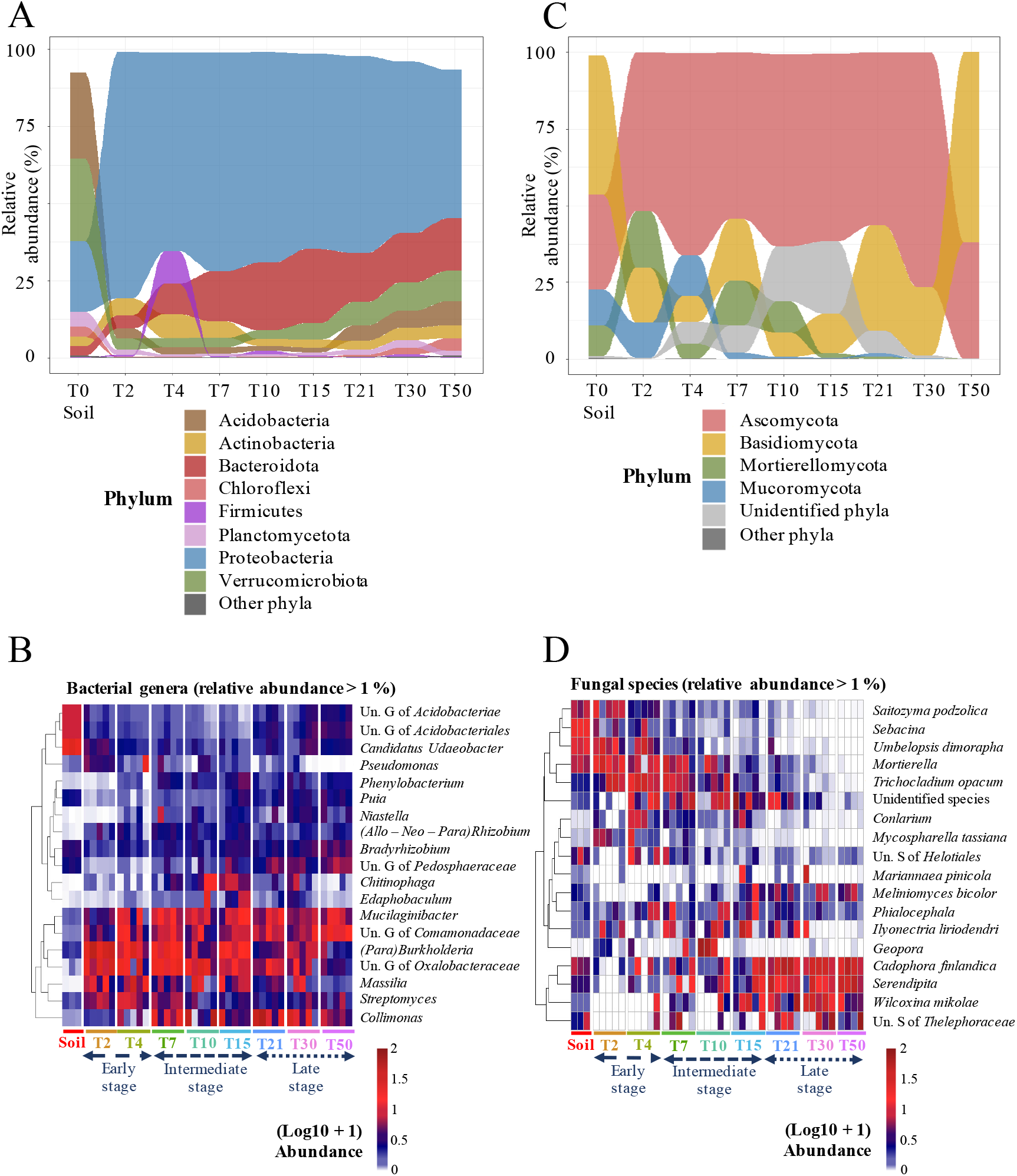
Composition of bacterial and fungal communities colonizing Populus roots across time. Average representation of the distribution of the most abundant bacterial (**A**) and fungal (**C**) phyla (> 2% in relative abundance) detected in bulk soil and in Populus roots collected at each sampling time from T2 to T50. Heat-map showing the relative abundance of bacterial genera (> 1% in relative abundance, **B**) and fungal species (> 1% in relative abundance, **D**) across compartments (bulk soil and roots) and sampling times (from T2 to T50). The cladogram on the left shows the similarity of the microbial taxa in terms of their relative abundance based on Ward’s minimum variance hierarchical clustering. Relative abundances were normalized through log(10) +1 transformation.

Fungal colonization of the roots was characterized by a significant expansion of Sordariomycetes (5.5x) to the expense of Agaricomycetes (−4.5x), Letiomycetes (−3.5x) and Pezizomycetes (−3x) compared to bulk soil samples (**Table S2**, P.adj <0.01). However, changes in fungal community composition of the roots were not significant at other taxa levels. As for bacteria, fungal communities detected in roots were dominated by a few species: nine fungal species accounted for 72% (± 3.4) (e.g. *Umbelopsis dimorpha*, *Trichocladium opacum* **Table S2**). However, the relative abundance of these dominant species was similar in the bulk soil and in roots at T2-T4, by contrast with bacteria. All dominant species at this stage except two (*Cortinarius* and *Sebacina*) were saprophytes or putative endophytes. As a consequence, fungal communities of roots were depleted in EcM (−5x) and endophytes (−3.5 x) compared to the bulk soil at the early stage (**Table S3**, P.adj <0.01). In particular, Thelephoraceae were absent from the root systems while *Sebacina* were found in low abundance compared to the bulk soil (2.1% ± 1.0, −8x).

### Evolution of the compositions of bacterial and fungal communities associated to *Populus* roots along time

The composition of root bacterial and fungal communities clearly evolved over time, from T2 to T50. The relative abundances of the eight most abundant bacterial phyla (> 1%) significantly varied from T2 to T50 (**Figure 3 A**, **Table S1**, Kruskal-Wallis, P.adj < 0.05). While still being the most dominant phylum, the proportion of Proteobacteria slowly decreased over time from 79.6% (± 2.8) at T2 to reach 48.6% (± 2.1) at T50 (**Figure 3 A**, **Table S1**, P.adj <0.05). Proteobacteria were replaced by members of the Bacteroidota, Verrucomicrobiota, Acidobacteriota and, in a lower extent, by Chloroflexi and Myxococota (**Figure 3 A**). All the genera that dominated in the roots in the early stage displayed decreased relative abundances over time and all were in the minority at T50. The only exception was the genus *Mucilaginibacter* that has remained among the dominant genera in roots along time. By contrast, the relative abundance of members of the *Chitinophagaceae*, in particular OTUs belonging to the genera *Chitinophaga*, *Edaphobaculum*, *Sphingomonas*, increased to reach a maximum of 12.9% (± 1.8) at the intermediate stage and then decreased to 5.5% (± 1.8). Finally, the relative abundances of *Comamonadaceae, Ktedonobacteracea* and *Pedosphaeraceae* families and the *Bradyrhizobium* and *Puia* genera significantly increased over time to reach a maximum at T50 (**Figure 3 B**, **Table S1**, Kruskal-Wallis, P<0.05).

Like bacteria, the composition of fungal communities in the roots deeply evolved from T2 to T50 although the dynamic slightly differed. The roots were mainly colonized by saprophytic fungi that dominated the soil assemblage until the end of the early stage, suggesting a later selection of fungal communities colonizing *Populus* roots than for bacterial communities. In addition, the fungal colonization was highly variable from one root system to another, particularly at the early time points. Nevertheless, as for bacteria, all dominant taxa at the early stage were fully replaced in the roots by other taxa along time. The relative abundance of Mucoromycota in roots dropped after T7 while Mortierellomycota remained abundant until T10 then almost disapeared (**Figure 3 C**, **Table S2**, Kruskal-Wallis, P.adj < 0.05). Meanwhile, the relative abundance of Basidiomycota slowly increased to become dominant at T50 (61.9 ± 9.2 %). This replacement between taxa was accompanied by a change in the ratio between saprophytes, EcM and endophytes in roots: saprotrophs, that represented 40 to 50% of the fungal species at the early stage (30 species, e.g. *Trichocladium opacum*, *Saitozyma podzolica*, *Umbelopsis dimorpha*, **Figure 3 D**) were rapidly replaced by EcM at the intermediate stage (20 species) while the proportion of endophytes continuously increased to reach 59% (± 5) in the late stage (6 species, **Figure 4 A**). Interestingly, successional replacements occurred within EcM and endophytes guilds. The EcM *Sebacina* and *Geopora* were detected in some roots systems at the early and intermediate type points, respectively but none of them persist over time while Thelephoraceae and *Wilcoxinia* gradually took over the root systems. Similarly, the endophytes *Leptodontidium* and *Phialocephala* were found only at the intermediate stage while *Serendipita* and *Cadophora finlandica,* that were already detected in low abundance at T2, significantly increased from intermediate to late stage representing respectively 27.0% (± 8.8) and 25.5% (± 8.9) of the relative abundance at T50 (**Figure 4 B**), and making them the major trophic group at T50 (52.5 ± 16.3%, **Table S3**).

**Figure 4.**
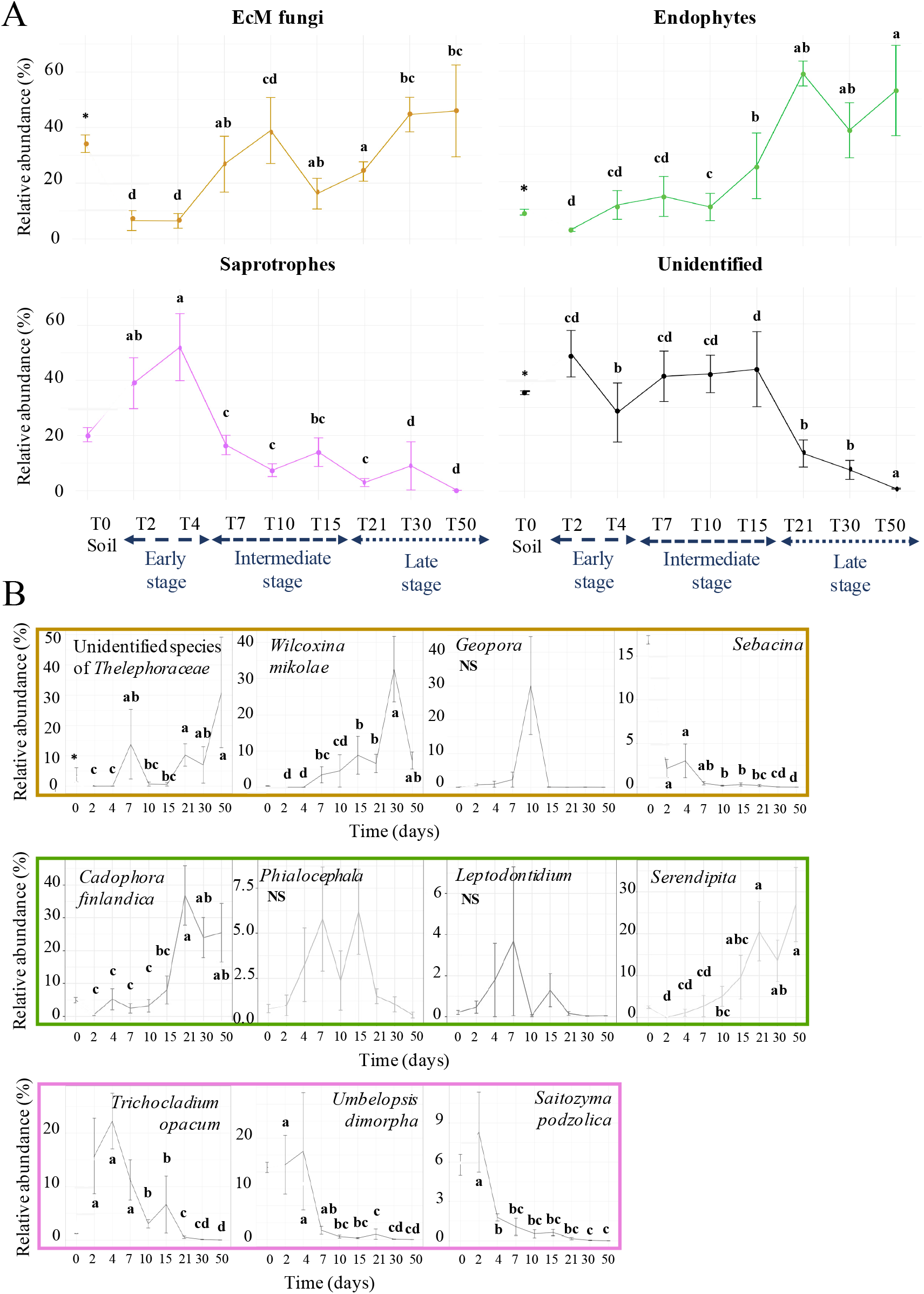
Evolution of fungal guilds in Populus roots along time. **A**. Relative abundance of the main fungal guilds detected in bulk soil and in Populus roots collected from T2 to T50 (3, 4 or 5 replicates +/− SE). **B**. Relative abundance of the most abundant EcM fungi (in yellow), saprotrophic fungi (in pink) and fungal endophytes (in green). The asterisks denote significant difference between bulk soil and Populus roots collected at T2 and different letters denote significant difference between each sampling time (Kruskal-Wallis, Benjamini and Hochberg correction, Fisher’s LSD post-hoc test, P.adj<0.05).

It is likely that some fungi and bacteria interact during the colonization of roots. To uncover such potential interactions, we looked for association patterns between the 13 fungal and 18 bacterial dominant families by sparse Partial Least Square Regression (sPLS). Strong associations were found between the relative abundances of Burkholderiaceae and Chaetomiaceae, and between Pedosphaeraceae and Heliotiales Incertae sedis and both were linearly correlated (Spearman p<0.01, **Figure 5**). In particular, the presence of unknown genera from the Pedosphaeraceae family (Verrucomicrobia) correlated with the one of the endophytic fungus *Cadaphora* (r^2^ = 0.68, p=2.e^−06^).

**Figure 5.**
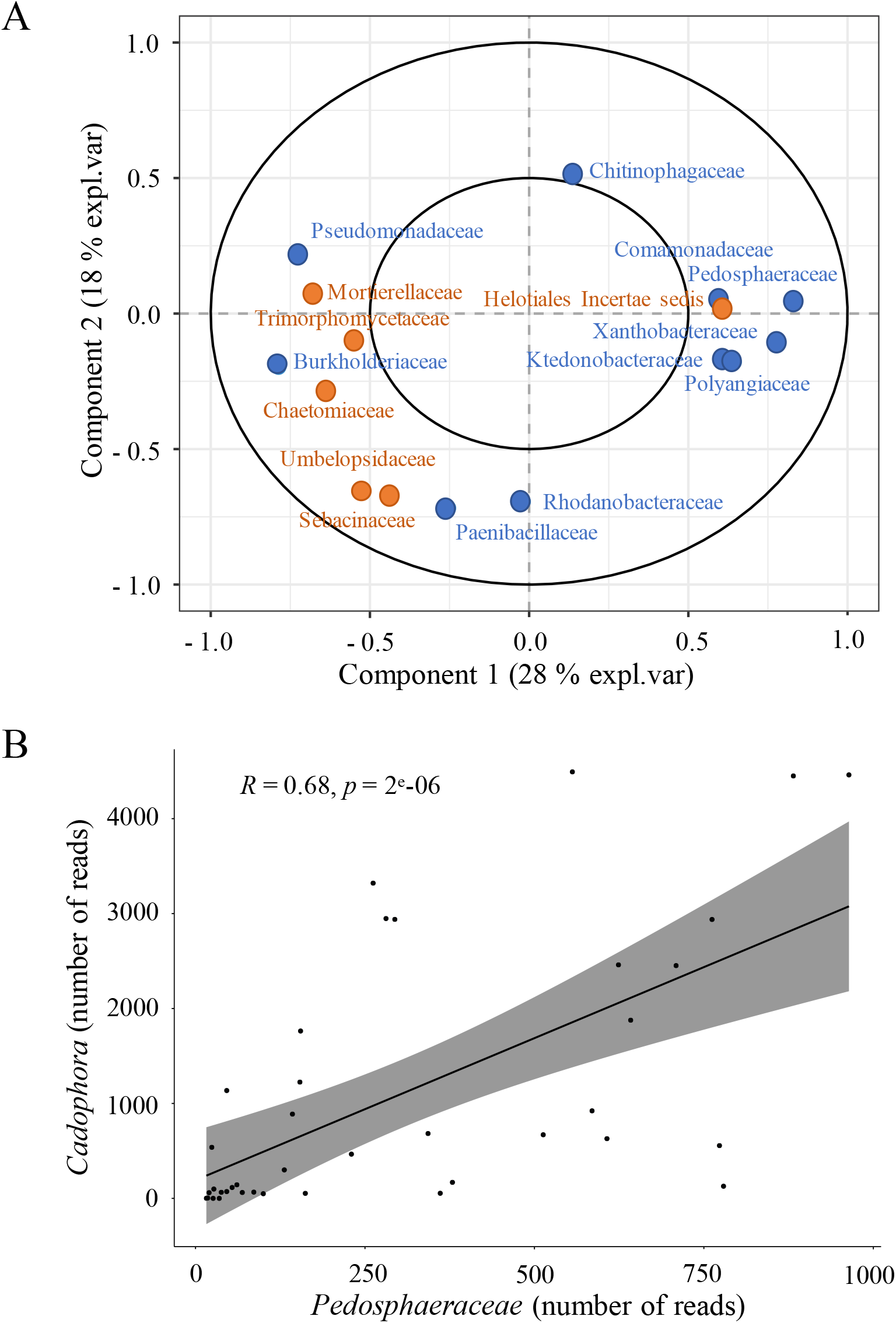
Association patterns between the most dominant microbial communities detected in roots along time. **A**. sparse Partial Least Square regression (sPLS) of 18 bacterial (in blue) and 13 fungal (in orange) dominant families detected in roots along time. **B.** Correlation between the number of reads of the endophyte fungi Cadophora and the number of reads of the unknown genus from the Pedosphaeraceae family (Spearman correlation test, P<0.01).

### Monitoring of fungal colonization in *Populus* roots by CLSM

MiSeq results brought global information about the structure and composition of microbial communities without knowledge about their spatial distribution and their physical interaction with the roots and with other microorganisms. In order to deepen our understanding of the process of root colonization and its dynamic, fungal colonization of the roots was also followed by CLSM.

As for high throughput sequencing, the first fungal presence was detected by CLSM between T2 and T4 (**Figure 6 A**). We observed spores and hyphae colonising the surface of root systems mainly from the apex (**Figure 6 B**). These colonisations were very heterogeneous from one sample to another, some root apexes being entirely surrounded by fungal mycelia while others had few hyphae (**Figure 6 C**). The fungal hyphae were septed with a diameter under 1μm and we observed a very low diversity of morphologies. After four days of growth, we observed the presence of dark septate endophytes (DSE) by light microscopy (**Figure S3**). Their hyphae were either extracellular or intercellular, but it was difficult to assess if they were intracellular or within apoplastic space.

**Figure 6.**
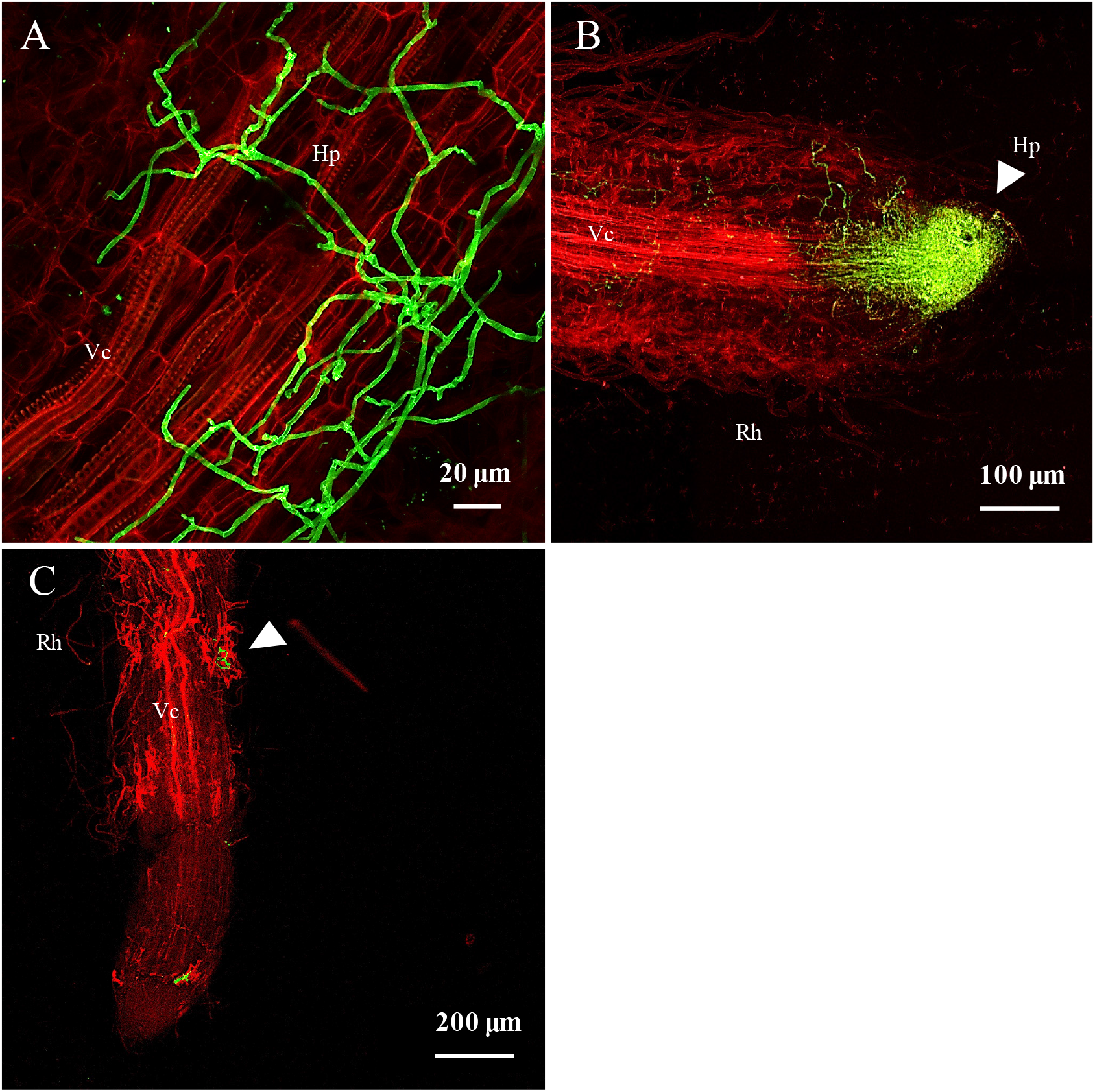
Early stage of the fungal colonization dynamic. **A**. Confocal microscopy images of poplar roots colonized by fungi after 4 days of culture. **A**. Extracellular hyphae surrounding a root after 4 days of culture. **B**. Hyphae accumulation at the apex of the root after 4 days of culture. **C**. Hyphae on root hairs after 4 days of culture. Fungal structures appear in green through WGA-Alexa Fluor 488 staining while root cell-walls were stained with propidium iodide and appear in red. Ap, Apex; Vc, Vascular cylinder; Hp, Hyphae; Rh, Root hair. Arrows indicate fungal hyphae colonization.

We detected an increased density of fungal morphologies by CLSM after 7 days. Fungal hyphae either developed between root cells, propagating in the apoplastic compartments particularly around epidermic regions (intercellular) or directly into root cells (intracellular) (**Figure 7 A, B**).

**Figure 7.**
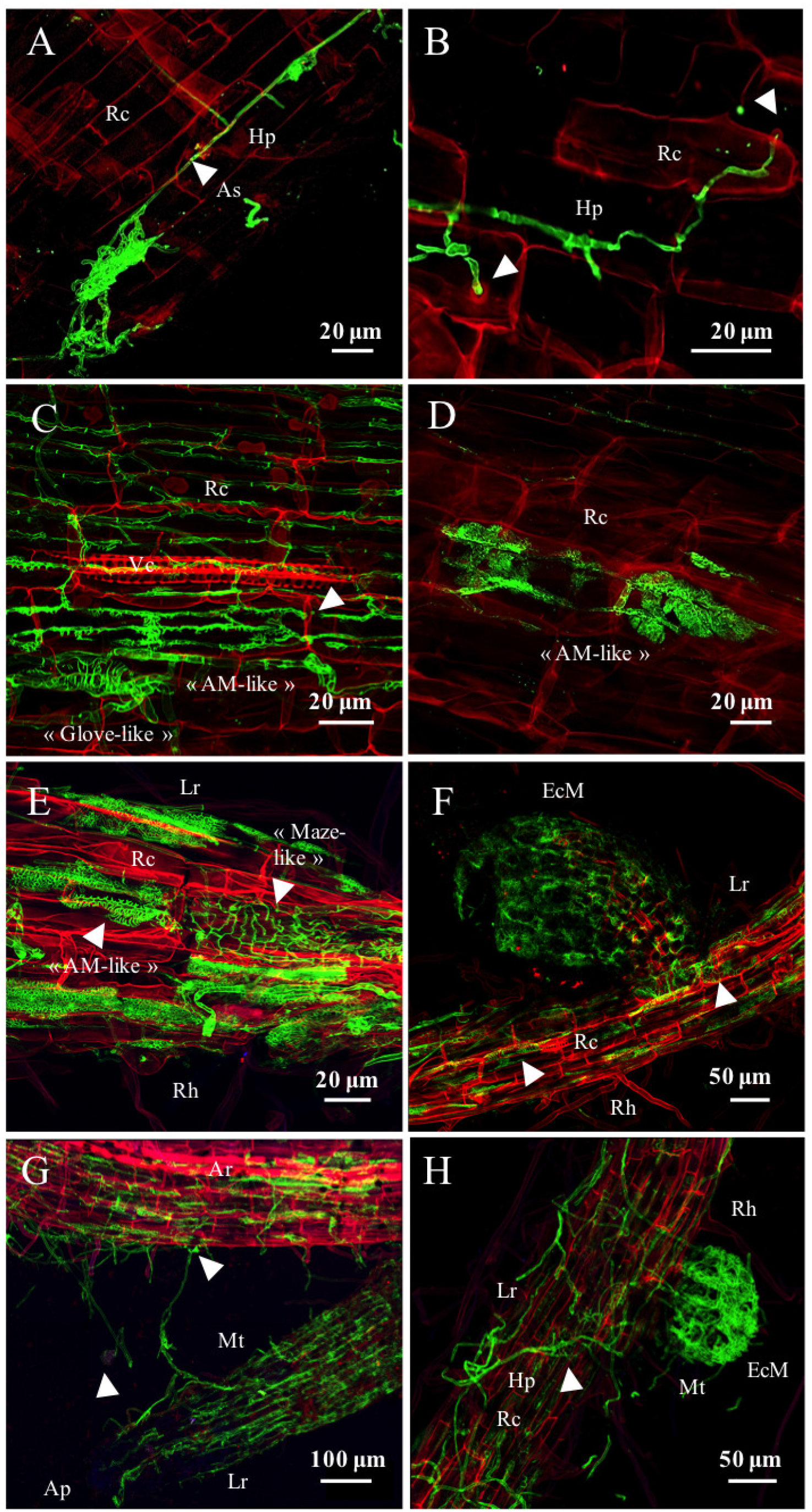
Intermediate and late stage of the fungal colonization dynamic. Confocal microscopy images of poplar roots colonized by fungi after 7 to 15 days (intermediate stage) and after 21 to 50 days (late stage) of culture. **A.** Development of fungal hyphae in the apoplastic space of cortical cells of poplar after 7 days of culture. **B.** Intracellular hyphal penetration in root cell after 7 days of culture. Arrows indicate the deformation of the root cell under the hyphal pressure. **C.** Co-existing fungal morphologies (**«** arbuscular like » and « glove-Hand like »), within the same root region after 15 days of culture. Arrows indicate hyphal intracellular penetration. **D. «** Arbuscular like » morphology observed in poplar root after 15 days of culture. **E.** At least three different morphologies are co-existing within the same root region after 21 days of culture. White arrow indicates the « maze like » structure and the « arbuscular like » structure. **F.** Mycorrhizal formation with co-existing fungal morphologies after 30 days of culture. White arrows indicate the « arbuscular-like » structure and the presence of the « maze like » fungal morphology, that seemed to be linked with the « glove-Hand like » morphology and the EcM forming structure. **G.** Hyphal propagation at T30 between the adventive root to the lateral root forming a probable EcM. White arrow indicates a germinated spore and the « hand glove like » morphology. **H.** EcM formation after 30 days of culture. Arrows indicate the « maze like » structure that seems to originate the EcM. Fungal structures appear in green through WGA-Alexa Fluor 488 staining while root cell-walls were stained with propidium iodide and appear in red. Ar, Adventive root; Ap, Apex; Hp, Hyphae; Lr, Lateral root; Mt, Mantle; Rc, Root cell; Rh, Root hair

The apoplastic colonisation stayed heterogeneous along the roots, and was dominantly present at the apex and in the root elongation zone. After 10 days, we observed an increase of both apoplastic and intracellular colonisation. Indeed, fungal hyphae were propagating from cell to cell by going through the root cell-walls and we were even able to see the pressure of the hyphae on the cell walls (**Figure 7 B**). Even though the global fungal diversity of morphologies remained poor at this stage of development, we noted the presence of septed and non-septed hyphae with diameters either inferior or superior to 1μm and we still observed the presence of DSE. After 15 days of culture, we observed an important increase of fungal density and morphological diversity in the root systems. We identified within the same root region the occurrence of distinct fungal morphologies with the dominance of two major structures (**Figure 7 C**). We detected the first dominant morphology in the intracellular compartment propagating from cell to cell and displaying an « arbuscular mycorrhizal-like » shape (**Figure 7 D**). The diameter of the hyphae was inferior to 1μm and hyphae developed by going through the cell walls from the epidermic to the central cells, forming a grid-shaped network. The second dominant morphology seemed to be propagating in both the intracellular and intercellular compartments, forming a “glove-like” shape with hyphae diameter closer to 5μm (**Figure 7 C**). This structure seemed to surround the root cell, reminding the Hartig net structure observed in ectomycorrhiza. The development of lateral roots after 15 days of culture was correlated with the establishment of the first distinct ectomycorrhizal structures (**Figure S3)**. Most EcM root tips already exhibited a mantle and a Hartig net (**Figure 7)**, however some EcM did not have a fully formed mantle and hyphal colonization seemed to originate from the adventive root system. The density of colonization and the occurrence of ectomycorrhizal structures were heterogeneous among the different root systems. Nevertheless, many fungal morphologies were present within the same region, both on lateral and adventive roots (**Figure S4**).

From 21 to 50 days of growth, we observed a global increase of the fungal density within the same root region with some roots systems being colonised from the apex to the top of the root at 50 days, even though it remained heterogeneous between the different root systems. We still observed DSE, both inter or intracellular and we detected two new abundant fungal morphologies that were sometimes located within the same root region. The first structure was developing in the intracellular compartment in both adventive and lateral roots, displaying a globular shape with hyphae diameter inferior to 1μm (**Figure 7 E**). The second morphology was only present in lateral and mycorrhized roots, with hyphae diameter superior to 1μm and displaying a « maze like » structure (**Figure 7 E**). Its location between the inter/intra compartments, as well as its origin were difficult to determine, but it is noteworthy that it was often associated and seemed to develop within ectomycorrhizal structures (**Figure 7 F**). In addition, we observed fungal structures developing between the adventive and the lateral root forming a potential EcM (**Figure 7 G**). The apex of the lateral root was not colonized by any fungal structure suggesting that the EcM forming originated from pre-existing fungal structures on the adventive root. We also detected the presence of germinating spores with emerging hyphae colonizing the root cells (**Figure 7 G**). We observed an increase of EcM establishment (**Figure 7 H**) and we assessed by mycorrhizal counts under CLSM that 37% (± 1) of lateral roots were forming ectomycorrhizal structures at the end of the experiment, even if their presence was also variable depending on the root systems.

Regarding the lateral roots, we observed the successional replacement of the « arbuscular like » structures to the benefit of the « glove-like » structures, surrounding the root cells and looking like the Hartig net and EcM. We did not observe this pattern in the adventive roots, where the « arbuscular like » morphologies continued to develop among the « glove-like » structures.

## Discussion

The establishment of the plant root microbiome is a dynamic process involving rich communities of microorganisms with distinct trophic modes and functional abilites (13). It relies on a complex set of interactions between the roots and microorganisms and between microorganisms themeselves. If studies of the colonisation dynamic of roots by bacteria and, in a lower extent by fungi, have been performed on several herbaceous plants and crops, few studies has been done so far to understand the early colonisation dynamic of tree roots by complex microbial communities (27). Here, we developed a microcosm experiment to grow axenic poplars in natural soil and to track the colonization of the root system by microorganisms. The transfer of plantlets from axenic conditions to the microcosm did not induce visible stress to plantlets as they grew normally and developped short roots and EcM symbiosis at same rates and timing than in other systems (34). In addition, we observed a rapid and dynamic colonisation of the root system by both fungi and bacteria. We were able to track both AM and EcM fungi, suggesting that our microscom allowed a normal development and colonization of the root system. To our knowledge, this is the first study investigating the primary steps of the spatio-temporal colonization of tree roots by fungi and bacteria.

### The dynamics of *Populus* root colonization differed between fungal and bacterial communities

Previous studies suggest that the root microbiome assemble from the surrounding soil in a two-step process. Rhizodeposition would first fuel a recruitment at the vicinity of the roots within few days and is followed by the entry inside the roots and a regulation of the community composition by the plant-microbial interactions (12; 35; 36). In accordance with this model, colonization of the poplar adventitious roots started within 2 days for both fungi and bacteria and the initial root microbial community gradually evolved over time. However, the degree of selection and the pattern of evolution differed greatly between bacterial and fungal communities. Root bacterial communities were already clearly different from the bulk soil only after two days, suggesting that a very early selection was operating for bacteria. In rice, first bacterial recruitment occurred around 24 hours (36). Sampling at earlier time points would be necessary to determine with more precision the exact timing and the very first steps of the bacterial colonization of the poplar roots. Bacterial diversity increased then stabilized around 15-21 days in a similar dynamic than for rice (36) but the community composition kept evolving until 50 days. Long-term evolution of short lateral root and EcM tip-bacterial communites was observed in pine for 24 weeks (27), indicating that the bacterial community may not have reached the equilibrium by 50 days in the present case. However, no real “climax” can be expected regarding the microbiome of tree roots for over more than a few months since root microbial communities evolve with seasons (8; 37) and the age of trees (38).

In contrast to bacteria, the root fungal community partially mirrored the one of the bulk soil at the early stage. It started to clearly diverge from the soil community at the intermediate stage, suggesting a later selection process leading to a reduction of the diversity of fungi in the roots. This is likely due to (i) the differences in the growth capacity of the three trophic guilds of fungi colonizing the roots and (ii) the molecular dialogue necessary for the establishment of AM, EcM and endophytes in roots. Indeed, if saprophytes and some endophytes (e.g. *Mortierella, Ilyonectria*) can rapidly develop on rich carbon source, EcM are generally slow-growers while AM require stimulation by plant strigolactone to develop (39). Endophytic fungi could directly interact with tree roots and promote host growth indirectly by manipulating the microbial community composition and functioning and manipulating host phytohormones (15). Thus both mycorhizal and endophytic types are less susceptible to colonize roots within few days even though reads of some EcM were already detected in low abundance at the early stage.

Altogether, our result suggest that bacteria and fungi react differently to tree selection factors or that the tree would select the bacterial and the fungal community through different processes. Bacteria and some fungi would be rapidly and strongly responsive to root exudates while other fungi would be sensitive to more specific signals (e.g. strigolactones, flavonoids…).

### Saprotrophs dominated early microbial communities but were counterselected over time

Proteobacteria and particularly *Burkholderiaceae* dominated the early root bacterial community. This is in accordance with previous studies that showed a significant enrichment of OTUs from Proteobacteria and from *Burkholderia* in roots of different tree species (6; 11; 27; 28; 40). Representative members of the *Burkholderiaceae* family have a high ability to develop on root exudates (12; 41; 42). Similarly, fungal community detected after two and four days of growth in natural soil were dominated by saprotrophs. These observations suggest that saprotrophs, even rare in soil, are the fastest colonizers of tree roots, certainly due to newly available carbon source from the plant and root exsudates (13). Endophytes such as *Mortierella* are also likely able to quickly grow on root exudates. Members of the *Mortierella* genus are commonly detected in soils of forests and poplar plantation (28; 30; 43). Although their ecological role is poorly understood, those fungi are characterized by their rapid growth when encoutening rich media (44). This dominance of saprotrophs among the microbial communities at early time points advocates for an important role of root exudates and particularly primary metabolites in the early colonization of the roots by microorganisms. In accordance with this hypothesis, fungal colonisation was limited to the surface of the roots and occurred mainly at the apex (**Figure 6 A**), the area where most of primary metabolites are exudated (45). Nevertheless, almost all taxa that dominated at the early stage, whether fungal or bacterial, were replaced over time. For instance, the relative abundance of members of the *Burkholderiaceae* family decreased for the benefit of other well-known tree root colonizers such as *Bradyrhizobium*, *Rhizobacter* or *Sphingomonas* (46). Several phenomena could explained such evolution of the microbial communities: competition between microorganisms, slow growth of the late comers, evolution of the composition of the root exudates, selection by the tree, cross-kingdom interactions between bacteria and fungi… Previous work on the annual grass *Avena fatua* showed that the dynamic of root exudate chemistry and the bacterial preferences for substrates drive the bacterial community assembly over the full season of growth (12; 13). Whether such mechanism also applies on a shorter period of time needs to be tested, as nothing is known so far regarding the dynamic of poplar root exudates during root development. Secondary metabolites could also play a role in the process. Association study between the level of salicylates in poplar and microbial community composition suggest that colonization of the rhizosphere by a number of bacterial taxa and fungi (e.g. Mortierellomycota) could be influenced by levels of salicylic acid, populin and tremuloidin (31). Whether these compounds participate in the early selection of the microbiome needs to be tested. Finally, one cannot exclude also that the bulk soil community that serve as a reservoir evolved along time although this is less likely based on our other experiment (47).

More unexpected is the very early detection reads corresponding to EcM in the roots, before short lateral roots start to develop. Among those, some such as *Sebacina* or *Laccaria* were present already at the early stage but did not further develop. Others were detected at T7 before the development of the short roots and were further able to form true ectomycorrhizae (*Thelephoraceae, Wilcoxina*). Similar early colonisation of primary root by EcM was found in *in vitro* experiments when inoculating eucalyptus roots with the ectomycorrhizal fungi *Pisolithus tinctorius* and *Paxillus involutus* (48) and *Betula pendula* with *Paxillus involutus* (49). Indeed, both studies found evidence of hyphal attachment to the roots after two days of inoculation with an accumulation of hyphae at the root apex. These observations would suggest that EcM can colonize the adventive primary roots before the formation of short lateral roots. It should be checked whether this step favours the EcM formation and requires a specific molecular dialog between the fungi and the plant as for the establishment of the symbiosis.

### The dominance of fungal saprotrophs vs EcM and endophytes fungi was reversed over time in *Populus* roots

The relative abundance of saprotrophic fungi dropped after 4 days in contrast with the relative abundance of EcM and endophytes which increased during the intermediate and late stages of root colonisation, reaching together 99% in relative abundance at the end of the experiment according to metabarcoding data (**Figure 4 A**). CLSM also revealed the abundant presence of AM fungi at the intermediate and late stages while AM fungal reads were almost completely absent from metabarcoding data (**Table S3**). Analysis of Illumina Miseq raw data before rarefaction indicated that reads corresponding to 10 OTUs belonging to the genera *Rhizophagus, Funneliformis* and the Archeosporales and Paraglomales orders could be detected in the roots but at very low levels that do not reflect the colonization of roots by AM as observed by CLSM. Such artefact and difficulty to properly track AM fungi in poplar roots by high throughput sequencing has already been highlighted by several authors (43; 50; 51) who suggest the use of additional methods for the analysis of AM interactions with poplar roots. Nevertheless, the combination of metabarcoding data and microscopy observations made it possible to draw the sequential events of the colonization of naïve poplar roots by fungi from the soil. Upon secretion of exudates in the rhizosphere, saprophytic fungi massively grew at the apices and on the surface of the roots for a few days. Whether plant defense mechanisms or interference competition with mycorhizal fungi and/or endophytes (52) put an end to their development needs to be determined. Meanwhile, endophytes including DSEs, AMs and EcMs slowly developed on the surface and started colonizing the inner tissues between T4 (DSE, potentially *Phialocephala*, *Cadophora* and *Leptodontidium*) and T7 (AM, EcM, other endophytes, potentially. *Serendipita*). At 10 days, first short roots formed and were colonized by at least AMs, DSEs and EcMs. However, if AMs continued to be present in the adventive roots they did not maintain themselves over time in short roots. Replacement of AM by EcM has been previously documented in eucalyptus (24; 25; 53) and poplar (26). However, these studies have been done on a long time scale, from 5 months to years, looking at the fungal colonisation dynamic of already grown-up trees and the mechanisms involved in such process are unknown.

At 30 days, inner tissues of adventitious roots were massively colonized but by fewer species than at earlier time points, mainly by the endophytes *Serendipita, Ilyonectria* and *Cadaphora* while EcM from the Thelephoraceae and *Wilcoxinia* likely developped functional ectomycorrhizae with Hartig net (**Figure 3 D**, **Figure 7 F-H**). As reported by (54), we noted by microscopy that some DSE were often associated with EcM. In addition to DSE, we also observed within a single ectomycorrhizal structures a diverse range of fungal morphologies, as also described recently in eucalyptus EcMs (55). These observations suggest that ectomycorrhizae in natural settings are made of more complex communites than “only” the root, the ectomycorrhizal fungus and associated bacteria; it would also include several additional endophytes whose relationship with the rest of the root-microbial community remains to be deciphered.

Fungal community compositions at T30 and T50 were very close, suggesting that an equilibrium may have been reached. However, this equilbrium would be only provisory as it is known that fungal community of trees evolves with seasons (8; 56) and all along their lifespan (57).

Interestingly, we observed the successional turn-over of distinct EcM and endophytic fungi in poplar roots. In both cases, fungal species that started to develop at the early to intermediate stages but that did not persist at the late stage are well-known members of the poplar root microbiome (e.g., *Mortierella*, *Umbelopsis*, *Sebacina*), suggesting that their exclusion is not due to a defensive reaction of the plant (15; 28; 51). If competition abilities of endophytes has not been investigated so far, (15) showed that a single endophyte species can shift the whole community of root-associated microorganisms. In addition, competition and priority effects in EcM fungi has been deeply scrutinized in the past (58). Both mechanisms are considered to have an important role in structuring communities of EcM. In the present case, EcM species that dominated in bulk soil (i.e *Sebacina*) and in the early time point did not took over the root system suggesting that priority effect was not the main process involved in the successional events of colonization. Yet, the important variability in the colonization of root systems by fungi at the early time point may result from stochastic events and priority effects. As *Thelephora terrestris* and *Wilcoxina* are considered to be highly competitive species (59; 60), we hypothesize that competitive exclusion is likely involved in the process. Potential interactions with other members of the microbiome may also be at play.

## Conclusions

In conclusion, this work has demonstrated that bacterial and fungal communities of the bulk soil successively colonized *Populus* roots. This colonisation took place in three major stages. The early stage was characterized by a massive colonization of the naïve roots by Proteobacteria members and saprotrophic fungi while the intermediate and late stages were charaterized by an increase of Bacteroidota, Verrucomicrobiota, Acidobacteriota (even if Proteobacteria still dominated bacterial communities), as well as of endophytic and EcM fungi. The establishment of root bacterial communities was stable earlier than fungal communities suggesting different establishment process between bacteria and fungi. Our observations constitute a first phase of exploration of the establishment of tree-microbes interactions as soon as roots appear and come into contact with the bulk soil. Future experiments should investigate the mechanisms involved in the formation of the root microbiome to disentangle the relative contributions of root exudates, plant defence and competition among microorganisms in this process.

## Material & Methods

### Biological material and sample preparation

*Populus tremula x alba* (INRAE clone 717-1B4) vitroplants were cultivated on Musharige & Skood (MS) supplemented with IBA (2ml.L^−1^) during one week before transfering them on MS for two weeks at 24°C in growth chamber (photoperiodicity of 16h, light intensity of 150 umol.m^−2^.s^−1^) until root systems were developed, as described in (61). Soil was collected from an 18-year-old poplar stand planted with *Populus trichocarpa* x *deltoides* and located in Champenoux, France (48° 51’ 46’’ N/2° 17’ 15’’ E). The first soil horizon (0-15 cm) was collected over an area of about 1m^2^ (i.e. approximately 50 kg of soil) and after pruning of brambles and adventitious plants and litter removal with a ratle. Then, soil was maintained at room temperature and homogenised through sifting at 2mm and fixed at 75% of humidity. Fifty grams of bulk soil was sampled in triplicate and stored at −20°C until DNA extraction.

Rooted vitroplants were selected to be homogeneous in terms of the size of the aerial part and the root system (i.e. approximately 1cm long for aerial parts and 2cm for roots). Selected vitroplants were transplanted in natural soil in transparent plastic pots with a filtered cover allowing gas exchange and a dark area at the ground level to prevent algae development. Plants were cultivated in a growth chamber (photoperiodicity of 16h, light intensity of 150 umol.m^−2^.s^−1^). Humidity in pots was maintained at 75% during all the experiment by weigthing pots and regular watering. Vitroplants were harvested after 0, 2, 4, 7, 10, 15, 21, 30 and 50 days of growth (**Figure S1**). At the beginning of the experiment (time-point “T0”) and at each time point, the root system of five plants were harvested, rinsed with sterile water, placed next to a ruler, photographed (Nikon Coolpix P530), freezed in liquid nitrogen and stored at −20°C until DNA extraction. Two additional plants were harvested and roots fixed in a solution containing 1 volume of 1X phosphate-buffered saline (PBS: 0,13 M NaCl, 7 mM Na2HPO4, 3 mM NaH2PO4, pH 7,2) for 3 volumes of 3% para-formaldehyde (PFA) overnight at 4°C (62). At T30 and T50 time point, the root system was sufficiently developed to be splited in two equal part to performed these two technical approach on all plants.

### Monitoring of vitroplant growth and EcM root colonization monitoring

Total area of root systems were measured for each vitroplant collected at the different time points on scan images using ImageJ (63) before freezing in liquid nitrogen or PFA fixation. Mycorhization rate of each vitroplants was quantified as previously described (64). Briefly, each root system was observed under a dissecting microscope. For each root system, 100 short roots were randomly examined and assessed as mycorrhizal or non-mycorrhizal. Mycorrhization rate is defined as the number of mycorrhizal roots observed divided by the total number of short roots examined.

### Confocal laser scanning microscopy

Staining procedures of root systems and fungi were adapted from (65) protocol. In brief, fixed root systems were washed 3 times in one volume of 1X PBS and a last wash in 1 volume PBS / 1 volume of 96% ethanol before clearing them during 2h at 90°C in 20% KOH. After 3 washes in distilled water, samples were incubated overnight in 1X PBS containing 10 μg.ml^−1^ WGA-Alexa fluor 488 (ThermoFisher Scientific, Waltham, USA), a specific marker of the chitin of fungal cell walls. Then, root systems were washed in 1X PBS and incubated for 15 min in 1X PBS containing 10 μg.ml^−1^ of propidium iodide (a DNA intercaling agent that is excluded by intact cell membranes and stains plant walls regardless of cells viability (66)), before 3 washes in 1X PBS. Samples were mounted between slide and cover slip with a drop of SlowFade solution (Life Technologies). All root samples were observed with a ZEISS LSM 780 (ZEISS International) confocal laser scanning microscope (CLSM). WGA-AF488 was excited using 488 nm excitation wavelenght and detected at 500-540 nm whereas 561 nm excitation wavelenght and detection at 580-660 nm were used regarding propidium iodide. Maximum intensity projections were performed using the ZEN software with z-stack of 30 to 50 μm of width.

### Staining for optic microscopy and observation

Blue staining of fungal stuctures was adapted from (65) and (67). Cleared roots were incubated at 90°C in KOH 10% during 20 min. After few washes in distilled water, root systems were incubated for 10 min in 0.1 N HCl at room temperature. We removed HCl without washing and we incubated the root systems during 30 min at 90°C in acidified ink (5% Waterman ink, 20% lactic acid, 75% water). Finally, roots were washed in distilled water before being mounted between slide and cover slip with a drop of glycerol 20% for observation under the OLYMPUS BX41 optic microscope (JENOPTIK Progres Gryphax camera, AxioVision v. 4.8.2 sofware).

The presence of dark septate endophytes (DSE) by light microscopy is characterized by melanised septed hyphae that were not stained by WGA-Alexa fluor 488.

### DNA extraction, Illumina Miseq amplicon sequencing and quantification of microorganisms on roots

Approximatively 250 mg of bulk soil samples was used for each individual soil DNA extraction. Soil DNA was extracted using the DNeasyPowerSoil Kit following the protocol provided by the manufacturer (Quiagen, Venlo, the Netherlands). The root system of each vitroplant was crushed in liquid nitrogen with mortar and pestle and fifty mg of root tissue were used to extract DNA using the DNAeasy Powerplant Kit (Quiagen, Venlo, the Netherlands). DNA of all extractions were quantified with a Nanodrop 1000 spectrophotometer (Nanodrop Products, Wilmington, DE, USA).

A two-step PCR approach was performed to barcode tag templates with frameshifting nucleotide primers as described by (11). Forward and reverse primer mixtures were used to maximize phylogenetic coverage of bacteria and fungi. Primer mixtures for tagging bacterial amplicons were composed of 4 forwards and 2 reverses 515F and 806R primers screening the 16S rRNA V4 gene region in equal concentration (0.1μM; 47). Primer mixtures for tagging fungal amplicons were composed of 6 forward and 1 reverse for ITS2 rRNA region at equal concentration (0.1μM; 47). To inhibit plant material amplification, a mixture of peptide nucleotide acid (PNA) blockers targeted plant mitochondrial and chloroplast 16S rRNA genes and plant ITS nuclear rRNA gene were added in PCR reaction mixes (47). Polymerase chain reaction (PCR) were performed for three replicates of each sample (2μl isolated DNA at about 10 ng/μl) using 2.5x Phusion flash high fidelity master mix (ThermoScientific) with 1.5μl of forward and reverse primer mix, 0.75μl of PNA probe (5 nM) and 8.5μl of 0.2μm filtered UV treated DNA free water (Carl Roth, France) in a total reaction volume of 30μl per sample. Thermal cycler conditions for the primary PCRs for bacterial amplification in bulk soil and root samples were 30 cycles of 98°C for 5s, 78°C for 10s, 52°C for 20s and 72°C for 15s. Primary PCR condition for fungal amplification in bulk soil and root samples were 30 cycles of 98°C for 5s, 78°C for 10s, 55°C for 20s and 72°C for 15s. PCR products without addition of microbial DNA (negative control) and mock communities of known fungal or bacterial compositions were added as quality controls. Samples of 50μl (30 ng DNA per μl) were sent for tagging and MiSeq Illumina Next Generation Sequencing using 2×250 pair-end SOP to GeT PlaGe INRAE sequencing platform (Toulouse, France). The raw data were deposited in the NCBI Sequence Read Archive website (http://www.ncbi.nlm.nih.gov/sra) under the SRA study accession numbers SRR12474095-SRR12474100 for 16S data and SRR12474163-SRR12474204 for ITS data (Project PRJNA657694).

### Sequence processing

Bacterial and fungal raw sequences were further processed with FROGS (Find Rapidly OTU with Galaxy Solution; 68) implemented on the Galaxy analysis platform (69). Sequences were demultiplexed, dereplicated, sequence quality was checked, oligonucleotides, linker, pads and barcodes were removed from sequences. Then sequences were removed from data set, if they were non-barcoded, exhibited ambiguous bases or did not match expectations in amplicon size, meaning 50 to 700 nucleotides for fungal and 280 to 500 for bacterial sequences. Remaining sequences were clustered into operational taxonomic units (OTUs) based on the iterative Swarm algorithm, then chimeras and phiX contaminants were removed. OTUs with a minimum number of reads above 5.10^−5^ percent of total abundance were kept for further analyses for both bacterial and fungal OTUs as proposed by (68). Fungal sequences were further process using the ITSx filter implemented in FROGS, in order to discard sequences where ITS region has not been detected. Bacterial double affiliation was performed by blasting OTUs against SILVA database (70) whereas Unite fungi database (71) was used for fungal double affiliation. OTUs with blast identity < 85% for bacterial and < 80% for fungal were considered as chimera and were removed from the dataset. Blast identity percentage was considered lower for fungal affiliation in order to keep combined sequences allowed by FROGS. Finally, OTUs corresponding to chloroplasts or mitochondria were removed from the data set. For both fungal and bacterial data, per-sample rarefaction curves were produced to assess sampling completeness, using function *rarecurve()* in package Vegan v3.5-1 (72) in R (version 3.4.3; 73). Samples with insufficient number of sequences according to rarefaction curves were removed.

Based on these, subsequent analyses of diversity and community structure were performed on datasets where samples had been rarefied with the Phyloseq (74) package to achieve equal read numbers according to the minimum number of total reads in any sample (8,245 reads for fungi and 13,980 reads for bacteria). Microbial community composition and structure in bulk soil and roots data were further analysed by using Phyloseq package (74).

FUNGuild (75) was used to classify each fungal OTU into an ecological guild. OTUs identified to a guild with a confidence ranking to “highly probable” or “probable” were conserved in our analysis, whereas those ranking to “possible” or with multiple assignation were called “unidentified”. In this study, we focused on the evolution of the distribution of saprotrophs, endophytes and EcM fungi by averaging their relative abundance (± SE) between each biological replicate at the same time point. Other fungi types of fungi were classified as “unidentified”.

A special procedure was used for AM reads because they were lost in the cleaning process. Therefore, in order to extract reads from AM fungi, the same analysis was repeated with a threshold of minimum abundance of 1.10^−6^ percent and the data generated was only used for AM analysis.

### Statistical analyses

Statistical analyses and data representations were performed using R software (73, R studio v1.2.5001). Significant differences in the mycorrhization rate between root samples collected across time were detected by checking the normality of the data distribution with Shapiro-Wilk test followed by One-way ANOVA and Tukey HSD tests. Differences in fungal and bacterial community structures over time were tested using permutational multivariate analysis of variance (pairwise PERMANOVA) based on Binary distances and differences in structures were visualized using a non-metric dimensional scaling (NMDS) ordination, using Jaccard (based on presence/absence) and Bray-Curtis (condisidering OTU presence/absence as well as relative abundance of OTU) dissimilarity matrices. Kruskal-Wallis test followed by a Benjamini and Hochberg correction (fdr correction) with Fisher’s least significant difference (LSD) post-hoc tests were used to detect significant differences in the relative abundance of fungal and bacterial phyla, orders and genera of the bulk soil and across root systems collected at the different time points. This procedure was also used to compare relative abundance of fungal guilds between root samples collected at the different time points with the difference that Bonferroni correction was allplied instead of Benjamini and Hochberg. Variations in fungal and bacterial diversity and richness were tested using Kruskal-Wallis test followed by a Bonferroni correction with Fisher’s least significant difference (LSD) post-hoc tests. HeatMaps of taxonomic relative abundances were produced using the R package pheatmap (76), and cladograms were built based on Ward’s minimum variance hierarchical clustering. As well, alluvial plots representing the bacterial and fungal relative abundance were produced using the ggalluvial package in R (77). Sparse Partial Least Squares regression (sPLS) methodology were used to look for associations between bacterial and fungal communities (mixOmics package, 78). Correlations between the number of reads of specific microbial taxa were calculated by using Spearman correlation test.

## Acknowledgments

We wish to thank Milena Gonzalo (INRA Nancy) for her helping in cultivating poplar vitroplants.

## Supplemental figure legends

**Figure S1** - Experimental design and approach used in this study.

**Figure S2** - Confocal microscopy image of axenic Populus tremula x alba adventive root at the beginning of the experiment (T0). Plant cell walls were stained with propidium iodide and appear in red. Ar, Adventive Root; Vc, Vascular cylinder

**Figure S3** - Dark Septate Endophyte (DSE) colonizing poplar roots from 4 to 30 days of culture. Optic and confocal microscopy images of poplar roots colonized by DSE. **A.** External colonization of roots by DSE after 4 days of culture. **B.** Extracellular DSE hyphae surrounding an adventive root at 10 days of culture. **C.** Extracellular DSE hyphae after 21 days of culture. Arrow indicates the DSE septa. **D.** Intracellular DSE hyphae propagating in the apoplastic compartment after 21 days of culture. Arrow indicates the DSE septa. **E.** Extracellular DSE hyphae surrounding an EcM forming on a lateral root after 30 days of culture. Arrow indicated the DSE hyphae. **E, F.** Overlay of image and the green track in order to visualize both melanized and non-melanized fungal structures. Non-melanized fungal structures appear in green through WGA-Alexa Fluor 488 staining. Ar, Adventive root; Hp, Hyphae; Lr; Lateral root; Rh, Root hair.

**Figure S4** - Intermediate stage of the fungal colonisation dynamic. Confocal microscopy images of poplar roots colonised by fungi after 7 to 15 days of culture. **A.** EcM formation on a lateral root after 15 days of culture. **B.** Hartig net formation on EcM after 15 days of culture. **C.** Co-existing and abundant fungal morphologies within the same root region after 15 days of growth. Orange arrow indicates the « arbuscular-like » and white arrow indicates the « hand glove-like » fungal structures. Fungal structures appear in green through WGA-Alexa Fluor 488 staining while root cell-walls were stained with propidium iodide and appear in red. Ar, Adventive root; Ap, Apex; Hn, Hartig net; Hp, Hyphae; Lr; Lateral root, Mt, Mantle; Rc, Root cell; Vc, Vascular cylinder

## Supplemental Table legends

**Table S1 – Relative abundance of bacterial communities at each taxonomic rank detected in the bulk soil and/or roots samples collected from T2 to T50**. Each given value is the average value of 3, 4 or 5 replicates ± SE. The second column denote significant differences between bulk soil samples and root samples collected from T2 and the third column denote significant difference in relative abudance of fungal communities between each sampling time from T2 to T50. Different letters denote significant difference between each sampling time (Kruskal-Wallis, Benjamini and Hochberg correction, Fisher’s LSD post-hoc test, P.adj<0.05).

**Table S2 – Relative abundance of fungal communities at each taxonomic rank detected in the bulk soil and/or roots samples collected from T2 to T50**. Each given value is the average value of 3, 4 or 5 replicates ± SE. The second column denote significant differences between bulk soil samples and root samples collected from T2 and the third column denote significant difference in relative abudance of fungal communities between each sampling time from T2 to T50. Different letters denote significant difference between each sampling time (Kruskal-Wallis, Benjamini and Hochberg correction, Fisher’s LSD post-hoc test, P.adj<0.05).

**Table S3 – Distribution of the relative abundance of fungal guilds detected in the bulk soil samples and in the roots samples collected from T2 to T50**. Each given value is the average value of 3, 4 or 5 replicates ± SE. The second column denote significant differences between bulk soil samples and root samples collected from T2 and the third column denote significant difference in relative abudance of fungal communities between each sampling time from T2 to T50 (Kruskal-Wallis, Bonferroni correction, Fisher’s LSD post-hoc test, P.adj<0.05).

