## Supplemental Figures for "Colonization of naïve roots from *Populus tremula x alba* involves successive waves of fungi and bacteria with different trophic abilities"

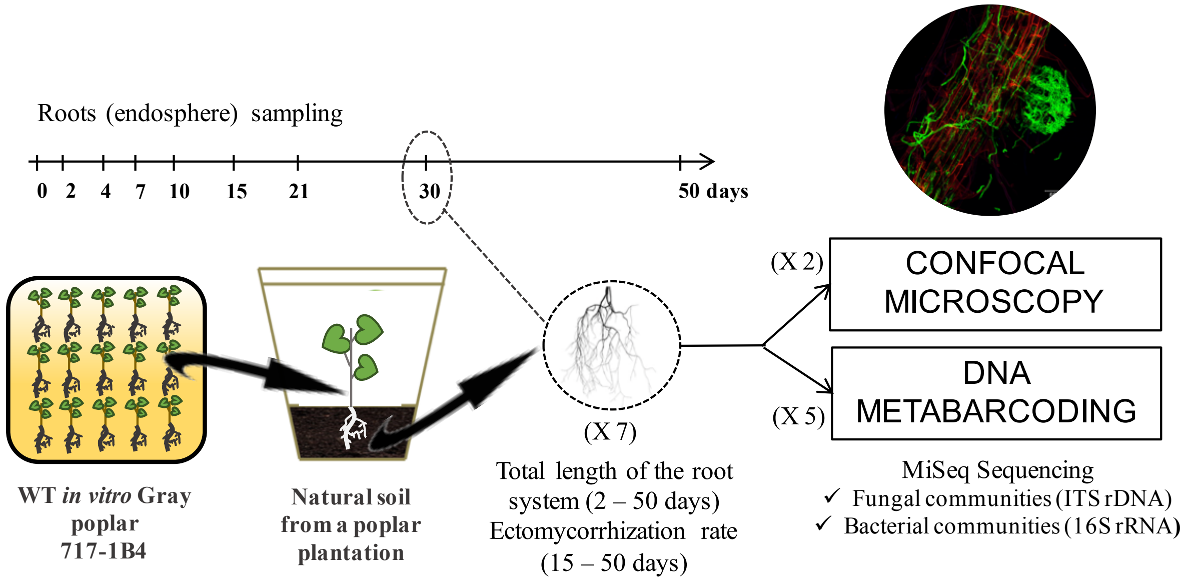


***Figure S1*** *- Experimental design and approach used in this study.*

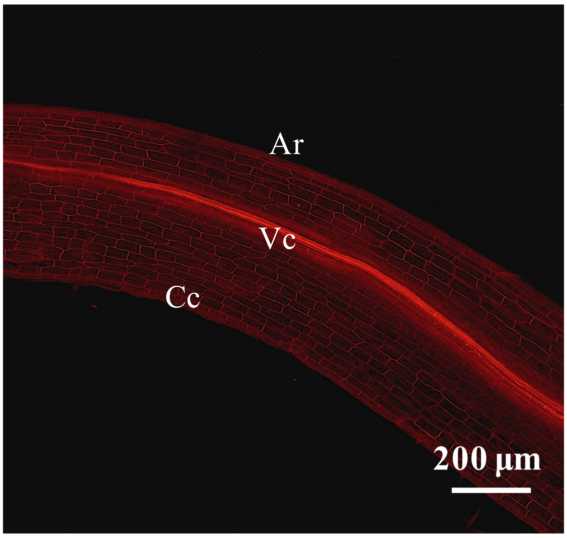


***Figure S2*** *- Confocal microscopy image of axenic Populus tremula x alba adventive root at the beginning of the experiment (T0). Plant cell walls were stained with propidium iodide and appear in red. Ar, Adventive Root; Vc, Vascular cylinder*

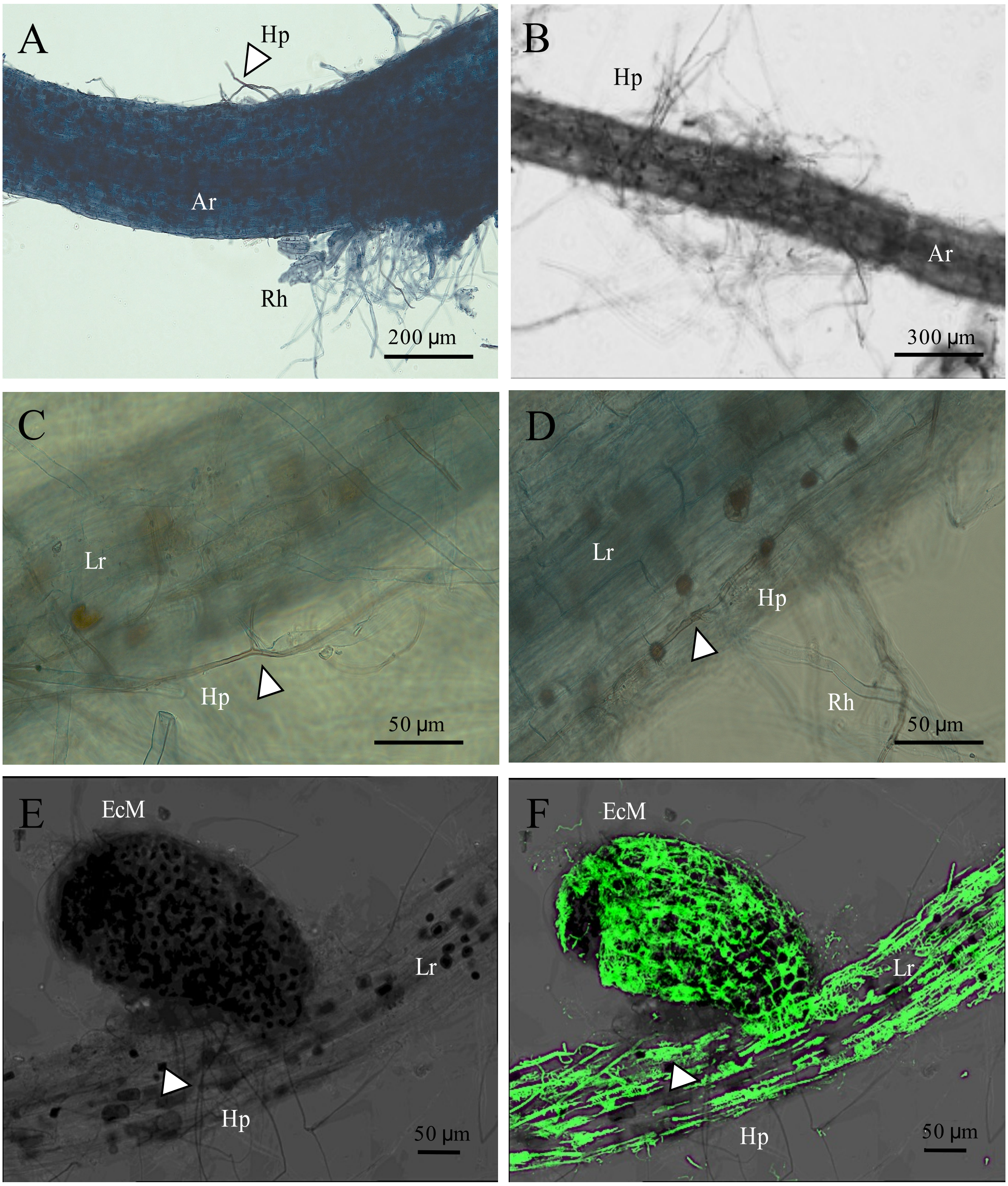


***Figure S3*** *- Dark Septate Endophyte (DSE) colonizing poplar roots from 4 to 30 days of culture. Optic and confocal microscopy images of poplar roots colonized by DSE.* ***A.*** *External colonization of roots by DSE after 4 days of culture.* ***B.*** *Extracellular DSE hyphae surrounding an adventive root at 10 days of culture.* ***C.*** *Extracellular DSE hyphae after 21 days of culture. Arrow indicates the DSE septa.* ***D.*** *Intracellular DSE hyphae propagating in the apoplastic compartment after 21 days of culture. Arrow indicates the DSE septa.* ***E.*** *Extracellular DSE hyphae surrounding an EcM forming on a lateral root*

*after 30 days of culture. Arrow indicated the DSE hyphae.* ***E, F.*** *Overlay of image and the green track in order to visualize both melanized and non-melanized fungal structures. Non-melanized fungal structures appear in green through WGA-Alexa Fluor 488 staining. Ar, Adventive root; Hp, Hyphae; Lr; Lateral root; Rh, Root hair.*

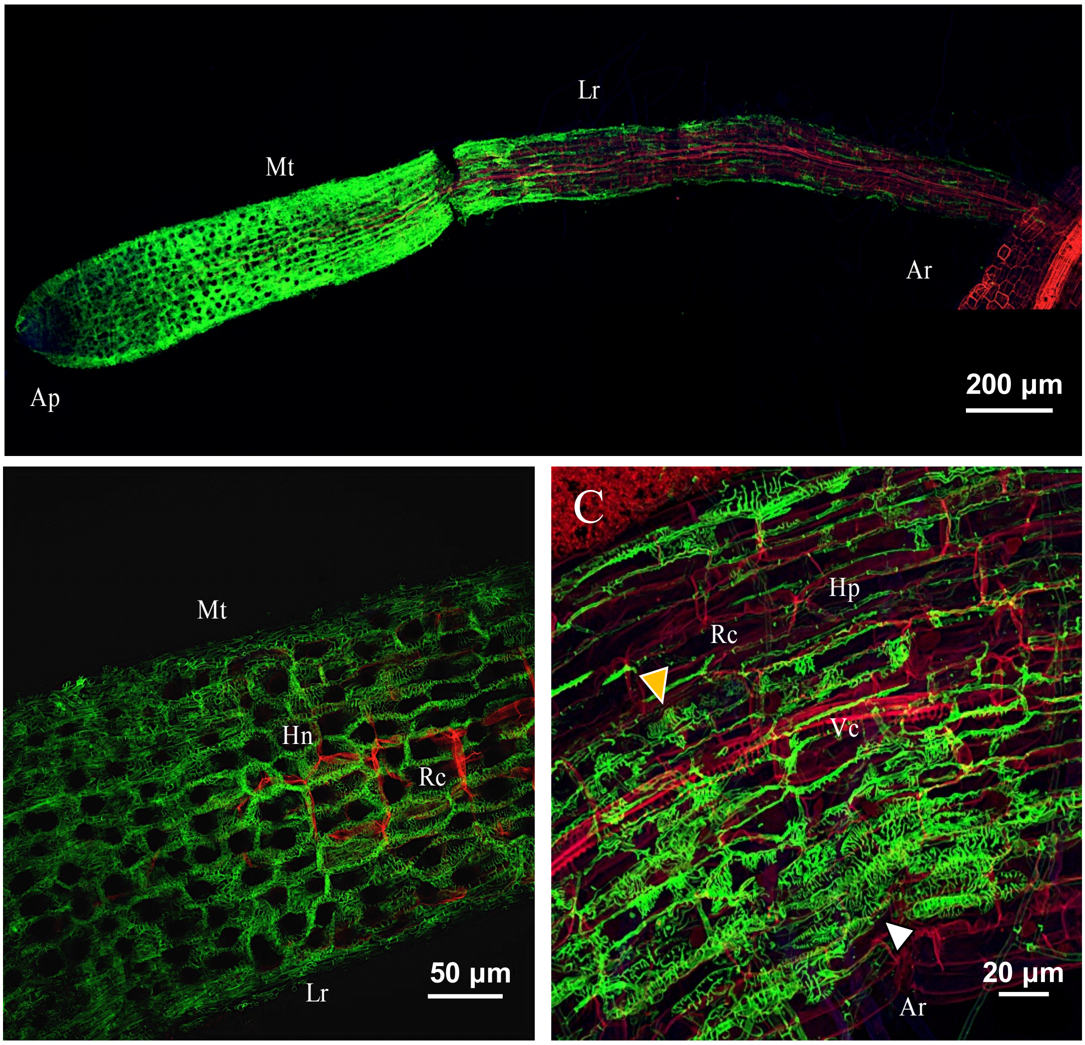


***Figure S4*** *- Intermediate stage of the fungal colonisation dynamic. Confocal microscopy images of poplar roots colonised by fungi after 7 to 15 days of culture.* ***A.*** *EcM formation on a lateral root after 15 days of culture.* ***B.*** *Hartig net formation on EcM after 15 days of culture.* ***C.*** *Co-existing and abundant fungal morphologies within the same root region after 15 days of growth. Orange arrow indicates the « arbuscular-like » and white arrow indicates the « hand glove-like » fungal structures. Fungal structures appear in green through WGA-Alexa Fluor 488 staining while root cell-walls were stained with propidium*

*iodide and appear in red. Ar, Adventive root; Ap, Apex; Hn, Hartig net; Hp, Hyphae; Lr; Lateral root, Mt, Mantle; Rc, Root cell; Vc, Vascular cylinder*
